# Engineering the Pseudomonas aeruginosa virus-isolation host viPAO1 reveals how host genetic barriers shape recoverable phage diversity

**DOI:** 10.64898/2026.09.16.752016

**Authors:** Anna Olina, Aleksei Agapov, Xinyan Yu, Rama P. Bhatia, MultiDefence consortium, Michael A. Brockhurst, Joanne L Fothergill, Stineke van Houte, Edze R. Westra

## Abstract

Cultured phage collections provide essential access to viral biology, but they capture only a small and biased fraction of natural phage diversity. Key barriers to diverse phage isolation are likely to include the prophages and defence systems present in the genomes of isolation hosts. Here, we show that removal of these host genetic barriers expands recoverable phage diversity in *Pseudomonas aeruginosa*. We constructed a series of progressively more permissive hosts derived from MPAO1 by sequentially deleting intracellular defence systems and resident prophages, culminating in viPAO1 (virus-isolation PAO1). Sequential removal of these barriers increased phage susceptibility, with Gabija and type I restriction-modification acting as major barriers to phage infection. In comparative isolation experiments, removal of intracellular barriers produced the largest increase in phage recovery, while additional receptor modification altered the composition of recovered phage subsets. viPAO1 also substantially increased recovery of phage contigs from environmental samples. Scaling this approach across 1090 clinical isolates enabled recovery of a curated collection of 69 individually purified temperate *P. aeruginosa* phages spanning 24 predicted genera, including 15 putative novel genera. Together, these findings demonstrate that removal of host genetic barriers to infection enhances recoverable phage diversity and establish permissive host engineering as a practical strategy for uncovering under-sampled regions of cultured phage diversity.

## Introduction

Bacteriophages (phages) are the most abundant biological entities on Earth and play central roles in shaping bacterial ecology, evolution and horizontal gene transfer [1–3]. They are increasingly recognised as valuable tools for biotechnology and as promising alternatives or complements to antibiotics for the treatment of multidrug-resistant bacterial infections [4–7]. Advances in metagenomic sequencing and computational identification of viral sequences have revealed an enormous reservoir of phage diversity in natural communities [8,9]. Yet comparison with experimentally isolated phages indicates that currently available cultured collections capture only a small and biased fraction of this diversity [8,9]. This disparity is particularly relevant for temperate phages, which are frequently identified as predicted prophage regions in bacterial genomes but are often not experimentally validated as infectious phage particles [10].

One contributor to this gap is bias inherent to phage isolation. Classical enrichment workflows and plaque assays preferentially recover phages that replicate efficiently under laboratory conditions, produce visible plaques and rapidly dominate mixed enrichments [11–13]. However, an equally important but comparatively underexplored source of bias is the bacterial host itself. Recovering a phage requires productive infection of the isolation host, making host genotype a biological filter that shapes which phages become experimentally recoverable.

Intracellular defence systems are likely to be a major component of this host-associated filter. Bacterial strains differ substantially in their defence repertoires, and these systems can restrict phage infection after genome entry, preventing replication of otherwise compatible phages. Phages unable to overcome such intracellular barriers may therefore remain experimentally undiscovered despite being present in the original sample. Supporting this idea, restoration of O-antigen synthesis together with removal of antiviral barriers in *Escherichia coli* K-12 enabled recovery of abundant environmental phage groups that were previously inaccessible using conventional laboratory hosts [14].

Resident prophages may provide an additional barrier to phage recovery. Prophages can restrict superinfection through mechanisms including repressor-mediated immunity, superinfection exclusion, surface modification and expression of additional defence functions [15,16]. Their presence may therefore further reduce the range of phages able to productively infect an isolation host.

In *Pseudomonas aeruginosa*, this problem may be especially pronounced. This species carries diverse and strain-specific repertoires of defence systems and prophages, both of which can influence phage infection. At the same time, many experimentally isolated *Pseudomonas* phage collections remain dominated by repeatedly recovered groups, suggesting that commonly used isolation hosts may favour particular phages while underrepresenting others [15,17–19]. We therefore hypothesised that engineering permissive *P. aeruginosa* hosts through removal of intracellular barriers would expand the diversity of phages that can be experimentally recovered.

To test this, we generated a series of progressively permissive MPAO1-derived hosts through sequential removal of resident prophages and intracellular defence systems, culminating in viPAO1 (virus-isolation PAO1), a highly permissive isolation host. We show that removal of intracellular barriers generates a gradient of host permissiveness and increases the recovery and diversity of experimentally accessible phages. We additionally tested whether modification of surface-associated susceptibility altered the composition of recovered phage populations. Together, our findings demonstrate that host genotype shapes experimentally recoverable phage diversity and establish permissive host engineering as a strategy for reducing isolation bias in *P. aeruginosa*.

## Results

### Construction of a panel of progressively permissive hosts

Candidate defence systems in *P. aeruginosa* MPAO1 were identified using DefenseFinder [11] and PADLOC [12]. This analysis identified candidate Gabija, Retron-I-B, type I restriction-modification and Helicase-DUF2290 defence systems. Resident prophages were predicted using PHASTEST [13] and revealed three predicted resident prophages, Pf4, a CTX-like prophage (prophage 2) and Pf6 (Fig. 1). We hypothesised that sequential removal of these intracellular defence systems and resident prophages would generate a panel of *Pseudomonas aeruginosa* hosts with progressively increasing permissiveness to phage infection. To test this, we constructed a series of MPAO1 derivatives in which intracellular defence systems and resident prophages were deleted sequentially, culminating in viPAO1, a strain lacking the Gabija, Retron-I-B, type I restriction-modification and Helicase-DUF2290 defence systems together with the resident prophages Pf4, CTX-like and Pf6 (Fig. 1A; Supplementary Tables 1 and 3c).

**Figure 1.**
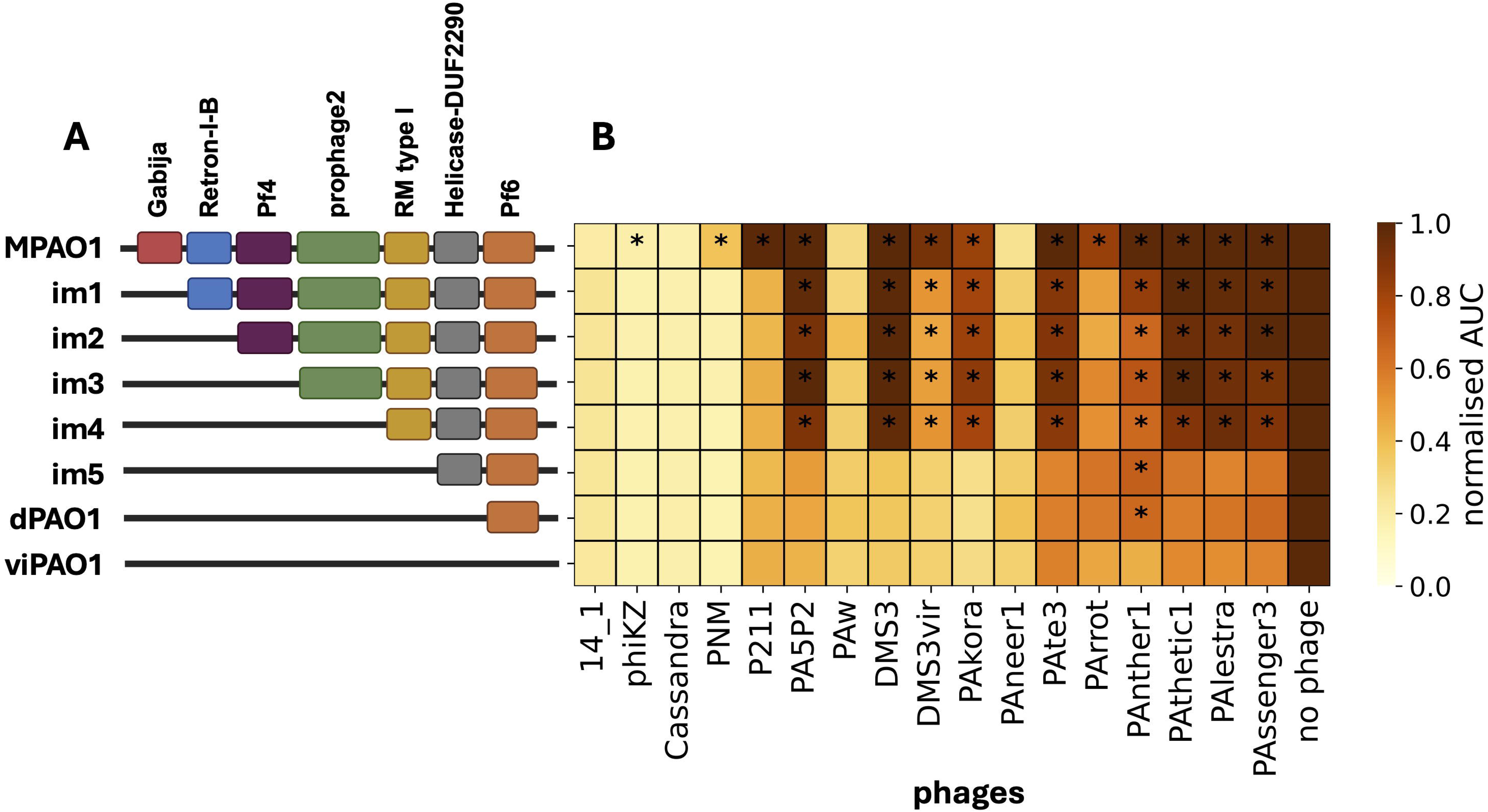
Sequential removal of intracellular barriers increases phage susceptibility. (A) Stepwise construction of progressively permissive strains derived from *Pseudomonas aeruginosa* MPAO1. Defence systems and resident prophages were removed sequentially, generating intermediate mutants im1-im5 and dPAO1 and culminating in viPAO1. Deleted elements included the Gabija, Retron-I-B, type I restriction-modification and Helicase-DUF2290 defence systems and the resident prophages Pf4, CTX-like and Pf6. (B) Susceptibility of the MPAO1 deletion series to a representative subset of lytic and temperate phages measured using growth-based infection assays. Bacterial growth was monitored by OD600, and phage susceptibility was quantified as the area under the growth curve (AUC) between 3 and 10 h post-infection. Heatmap values show normalised AUC, with lower values indicating stronger phage-mediated growth inhibition and therefore greater susceptibility. Asterisks indicate strain–phage combinations with significantly higher normalised AUC than viPAO1 after Benjamini–Hochberg correction for multiple testing (adjusted *P* < 0.05). Full susceptibility datasets for the lytic and temperate phage panels are shown in Supplementary Fig. 1.

All deletions were confirmed by PCR and whole-genome sequencing. Under standard laboratory conditions, none of the engineered strains displayed significant differences in growth compared with the parental MPAO1 strain in LB medium at 37°C (Fig. S1C), indicating that sequential removal of intracellular barriers did not measurably affect bacterial fitness under these conditions.

To determine whether successive deletions generated gradually more permissive hosts, we challenged the parental strain, intermediate deletion mutants and viPAO1 with a panel of lytic and temperate *P. aeruginosa* phages using a liquid infection assay (Fig. 1B; Fig. S1A,B; Supplementary Tables 4 and 5). Sequential removal of intracellular barriers resulted in an overall increase in host susceptibility to phage infection, confirming that the engineered strains represent a gradient of permissiveness.

Deletion of Gabija produced a major increase in susceptibility across the phage panel, while subsequent removal of the type I restriction-modification system caused an additional further increase in susceptibility. Together, these two defence systems represented the largest intracellular barriers to productive phage infection in the MPAO1 background. In contrast, deletion of Retron-I-B, Helicase-DUF2290 and the resident prophages produced more phage-specific effects, although each successive deletion increased susceptibility for subsets of phages, consistent with multiple intracellular barriers collectively shaping phage infection.

The effect of host genotype differed markedly between the lytic and temperate phage panels. Phage susceptibility was quantified from bacterial growth during liquid infection assays, with lower normalized growth-curve AUC indicating stronger phage-mediated growth inhibition. Among the temperate phages, 40 of 43 (93%) produced significantly lower normalized AUC in viPAO1 than in at least one intermediate deletion mutant, consistent with a progressive increase in susceptibility following sequential removal of intracellular barriers (Fig. S1B). In contrast, this pattern was observed for only 12 of 67 (18%) lytic phages, indicating that host genotype had a more limited effect across most of the lytic panel (Fig. S1A). This difference likely reflects, at least in part, the isolation history of the phage panels: many lytic phages used here were originally isolated on PAO1 or other unmodified hosts, whereas the temperate phage collection was assembled using the MPAO1-derived hosts dPAO1 and viPAO1, which lack most identified defence systems and resident prophages.

Together, these results demonstrate that sequential removal of intracellular defence systems and resident prophages generates a gradient of host permissiveness, with Gabija and the type I restriction-modification system representing the largest barriers in this genetic background. This strain panel therefore provided an experimental framework to test how host permissiveness shapes the diversity of phages recovered during isolation.

### Host genotype reshapes experimentally recoverable phage diversity

Having established that sequential removal of intracellular barriers generated a gradient of host permissiveness, we next asked whether increased permissiveness alters the diversity of phages recovered during isolation. We define recovered phages here as phage genomes detected in sequencing pools generated from plaques isolated on each host background. To test this, we first performed a controlled comparative isolation experiment using four prophage-rich clinical *P. aeruginosa* isolates selected based on prophage predictions from whole-genome sequencing data (Fig. 2A; Supplementary Table 1). Prophages were induced using mitomycin C, and the resulting lysates were enriched and plated on six host backgrounds: MPAO1, viPAO1 and their T4P- and LPS-associated receptor-mutant derivatives.

**Figure 2.**
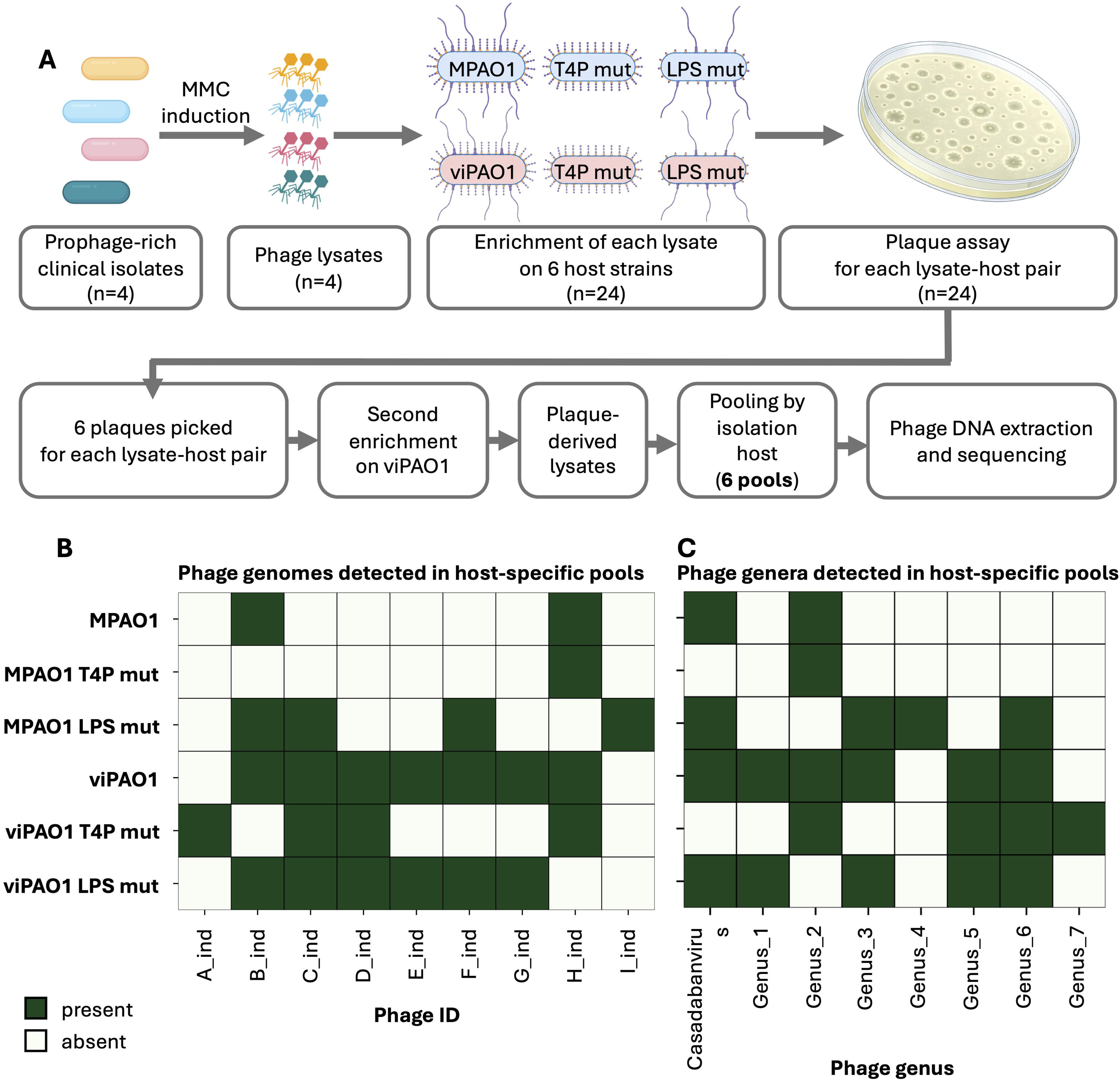
Host genotype shapes recovery of induced temperate phages. (A) Schematic overview of the comparative induced-phage isolation workflow. Prophages were induced from four prophage-rich clinical *P. aeruginosa* isolates using mitomycin C treatment. Induced lysates were enriched on six host backgrounds: MPAO1, viPAO1, and corresponding T4P- and LPS-associated receptor-mutant derivatives. Following plaque assays, six plaques were picked from each lysate-host pair. Plaque-derived lysates were secondarily enriched on viPAO1, then pooled by original isolation host background to generate six host-specific sequencing pools. Pooled phage DNA was extracted and sequenced using long-read sequencing. (B) Presence/absence matrix of induced temperate phage genomes detected in host-specific sequencing pools. Columns represent individual recovered phage genomes and rows represent the host backgrounds used for phage isolation. A phage genome or phage contig was considered present in a sample if primary alignments with MAPQ ≥20 covered at least 50% of its length or 10 kb, whichever was greater, and if at least five reads aligned to the genome or contig. (C) Genus-level summary of the phages detected in the host-specific sequencing pools. Genera were assigned using vConTACT3.

Removal of intracellular barriers caused a 3.5-fold increase in recovery of distinct temperate phages. Across the four induced lysates, MPAO1 recovered two phage genomes representing two genera, whereas viPAO1 recovered seven genomes representing six genera (Fig. 2B,C; Supplementary Table 6). Thus, increased permissiveness did not simply result in recovery of more closely related phage genomes but expanded both the number and taxonomic diversity of experimentally recoverable temperate phages.

We next asked whether receptor-associated mutations could further alter phage recovery. Because surface receptor composition is expected to influence which phages can infect and amplify on a given host, we compared MPAO1 and viPAO1 with their T4P- and LPS-associated receptor-mutant derivatives.

Receptor-associated mutations altered the composition of recovered phages in a host-background-dependent manner. In the MPAO1 background, the T4P-associated mutant recovered one phage genome representing one genus, whereas the LPS-associated mutant recovered four genomes representing four genera, compared with two genomes from two genera on parental MPAO1. In the viPAO1 background, the T4P- and LPS-associated mutants recovered four and six genomes, representing four and five genera, respectively, compared with seven genomes representing six genera on parental viPAO1. Thus, receptor modification altered the composition of recovered phage subsets but did not increase overall recovery beyond that achieved by removal of intracellular barriers.

To ask whether similar host-dependent recovery patterns extended beyond inducible temperate phages, we next performed exploratory environmental enrichments using four environmental samples: two wastewater samples, one biogas slurry sample and one activated sludge sample (Fig. S2; Supplementary Table 7). Because these samples contained mixed environmental phage communities, and because no matching bacterial source genomes were available to guide assembly as in the prophage induction experiment, the resulting assemblies were less complete and more difficult to resolve. We therefore refer to these recovered sequences as phage contigs rather than complete genomes. In addition, the presence of closely related environmental phages in pooled sequencing samples likely made this analysis conservative, with diversity more likely to be underestimated than overestimated.

Environmental enrichments showed a similar overall pattern. MPAO1 recovered one phage contig representing one genus, whereas viPAO1 recovered 16 contigs spanning three genera. Receptor modification again altered recovery. In the MPAO1 background, the T4P- and LPS-associated mutants recovered 16 and two contigs, representing two genera each, respectively. In the viPAO1 background, the T4P-associated mutant recovered 23 contigs spanning three genera, whereas the LPS-associated mutant recovered six contigs spanning two genera (Fig. S2). Thus, increased intracellular permissiveness enhanced environmental phage recovery, while receptor modification altered the composition of the recovered phage population.

These analyses showed that different host backgrounds recovered phages from multiple taxonomic groups. However, they did not show how the recovered phages were distributed relative to the broader genomic diversity of known *P. aeruginosa* phages. We therefore used vConTACT3 gene-sharing network analysis to compare induced-lysate phages and environmental phage contigs recovered in this study with available *P. aeruginosa* reference phages. Recovered phages and contigs were distributed across multiple regions of the reference network rather than being concentrated within a single gene-sharing cluster (Fig. S3). The 37 phage genomes and contigs recovered in the comparative isolation experiments spanned 8 of the 25 family-level groups and 12 of the 49 genus-level groups represented in the combined vConTACT3 network of recovered sequences and reference *P. aeruginosa* phages. Notably, the recovered sequences included one family-level group and seven genus-level groups that were not represented among the reference *P. aeruginosa* phages in the network. This indicates that host-dependent recovery extends across genomically distinct parts of *P. aeruginosa* phage diversity. Although the pooled datasets were less complete than the individually purified collection described below, the network analysis supports the conclusion that changing the isolation host altered access to multiple, broadly separated phage groups.

Together, our comparative isolation experiments demonstrate that host genotype strongly influences which phages become experimentally recoverable. Removal of intracellular barriers increased recovery of phage genomes and genera, while receptor-associated mutations altered the composition of recovered phage subsets. These results indicate that removal of intracellular barriers provides a broad increase in phage recovery, whereas receptor modification can redirect isolation towards distinct phage subsets.

### Permissive host backgrounds enable recovery of a broad curated temperate phage collection

Having shown that host genotype alters which phages are recovered during comparative isolation, we next asked whether combining permissive host backgrounds could be used at scale to recover a broad and taxonomically diverse temperate phage collection. We screened 1090 clinical *P. aeruginosa* isolates from diverse infection sites (Supplementary Table 2) for inducible temperate phages. Using viPAO1, dPAO1 and dPAO1 ΔpilZ as isolation hosts, we recovered 78 plaque-forming phage isolates. Of these, 75 formed stable lysogens under the conditions tested, whereas three did not form stable lysogens and were sequenced directly from phage lysates.

Because repeated induction and screening can recover the same phage more than once, we compared the 78 recovered isolates at the genome level. Two isolates were excluded because their genomes were identical to the previously described phages JBD69 and JBD24 [15,20]. Among the remaining newly recovered isolates, seven represented independent recoveries of phages with identical genome sequences and were collapsed to a single representative. This resulted in a curated non-redundant collection of 69 unique newly isolated phages (Supplementary Table 5). Independently recovered duplicate isolates were retained in the collection metadata and distinguished by numerical suffixes.

To improve genome completeness and annotation, phages were sequenced as lysogens in dPAO1 or dPAO1 dPilZ backgrounds rather than exclusively as free phage lysates (Fig. 3A). This approach enabled confident identification of prophage boundaries and chromosomal insertion sites, and reduced complications associated with direct sequencing and assembly of phage particles. Details for each phage isolate, including genome accession number, insertion site, known receptor information and isolation host, are provided in Supplementary Table 5.

**Figure 3.**
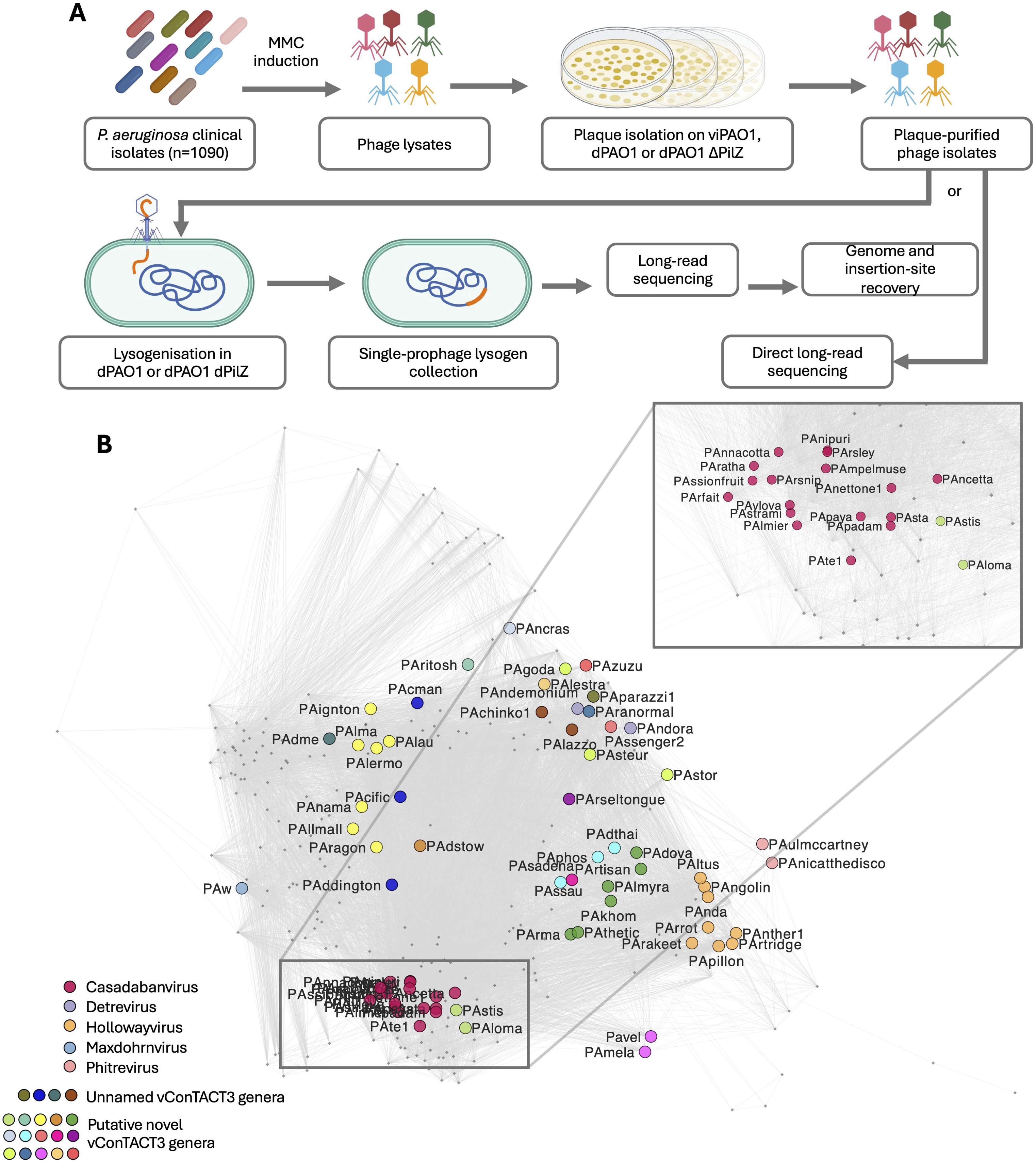
Permissive hosts enable recovery of a genomically diverse temperate phage collection. (A) Schematic overview of generation and genomic characterisation of the curated temperate phage collection. A total of 1090 *P. aeruginosa* isolates were screened following mitomycin C induction, and induced lysates were subjected to plaque isolation using viPAO1, dPAO1 or dPAO1 ΔpilZ. Seventy-eight plaque-forming phage isolates were recovered. Seventy-five formed stable lysogens in dPAO1 or dPAO1 ΔpilZ and were sequenced as lysogens using long-read whole-genome sequencing. Three phages isolated on viPAO1 did not form stable lysogens in these backgrounds under the conditions tested and were sequenced directly from phage lysates. Following exclusion of two isolates identical to the previously described phages JBD69 and JBD24 [15,20] and collapse of seven independent duplicate genome recoveries, 69 unique newly isolated temperate phages were retained for collection-level analyses. (B) vConTACT3 gene-sharing network comparing the 69 phages isolated in this study with publicly available *P. aeruginosa* reference phages. Nodes represent phage genomes and edges represent gene-sharing relationships inferred by vConTACT3. Phages isolated in this study are coloured according to their predicted genus, with each of the 24 predicted genera represented by a distinct colour; reference phages are shown in grey. The collection comprises 24 predicted genera, including 15 putative novel genera, and seven predicted families, including one putative novel family. The taxonomic composition and family-level genome-size distributions are shown in Supplementary Fig. 4.

Taxonomic analysis demonstrated that the collection captured substantial genomic diversity. The 69 temperate phages represented 24 predicted genera, including 15 putative novel genera, and seven families, including one putative novel family. The taxonomic composition of the collection and the distribution of genome sizes across predicted families are summarised in Supplementary Fig. 4. Thus, the use of permissive host backgrounds enabled recovery of a collection containing extensive previously under-sampled temperate phage diversity.

To place this collection in the context of known *P. aeruginosa* phage diversity, we performed vConTACT3 [16] gene-sharing network analysis together with publicly available *P. aeruginosa* reference phages (Fig. 3B). The phages isolated in this study were broadly distributed across the network rather than clustering within a small number of closely related groups. The 69-phage collection spanned 7 of the 23 family-level groups and 24 of the 59 genus-level groups represented in the combined vConTACT3 network of our isolates and reference *P. aeruginosa* phages. Some isolates occupied densely connected regions of the network, increasing representation within previously sampled parts of *P. aeruginosa* phage diversity space, whereas others were positioned towards network edges, consistent with lower gene sharing with available reference genomes. This pattern indicates that the collection both expands known phage diversity and increases sampling depth within existing groups.

Together, these results show that combining permissive and receptor-modified host backgrounds enables recovery of a broad, genomically diverse and taxonomically rich temperate phage collection, including numerous previously under-sampled lineages.

## Discussion

Experimentally recoverable phage diversity is shaped not only by the environment being sampled, but also by the biological properties of the host used to culture that diversity. In this study, we show that host-associated barriers strongly influence which *Pseudomonas aeruginosa* phages become experimentally recoverable during isolation. By sequentially removing intracellular defence systems and resident prophages from MPAO1, we generated viPAO1, a highly permissive isolation host, and showed that these deletions created a measurable gradient of phage susceptibility. Comparative isolation experiments further demonstrated that permissive and receptor-modified host backgrounds recover partially overlapping but distinct phage subsets. Finally, by combining complementary permissive hosts, we assembled a curated collection of 69 individually purified temperate phages spanning broad genomic and taxonomic diversity (Supplementary Table 5). Together, these findings establish host engineering as a strategy for reducing host-associated isolation bias and expanding access to under-sampled cultured phage diversity.

Our results support the growing view that phage isolation workflows are inherently selective. Conventional enrichment and plaque-based methods favour phages that replicate efficiently under laboratory conditions, dominate mixed cultures and produce visible plaques [14,17,18]. Recent efforts to develop plating-independent approaches, such as droplet-based isolation, highlight how many phages may remain inaccessible when recovery depends on visible plaque formation [18]. Similarly, large-scale analyses of cultured phage genomes show that available phage databases remain strongly biased towards particular host genera, phage lifestyles and repeatedly sampled groups [10,19]. Our findings extend this concept by showing that, even within a single bacterial species and from the same starting material, the genotype of the isolation host can strongly alter which phages are recovered.

Intracellular barriers were major determinants of host permissiveness in MPAO1. Among the deleted elements, Gabija and the type I restriction-modification system produced the clearest increases in phage susceptibility, indicating that these systems represent major filters to productive infection in this background. This is consistent with recent mechanistic work showing that Gabija is a widespread prokaryotic defence system that protects cells through DNA-targeting activity [21,22]. Our type I restriction-modification result is also consistent with recent work showing that deletion of the type I restriction endonuclease from PAO1 increases phage propagation and improves recovery of phages from environmental samples [23]. Importantly, however, Gabija and type I restriction-modification were not the only relevant barriers. Other defence systems and prophage deletions produced more phage-specific effects, consistent with a multilayered model in which different phages are restricted by different host-associated barriers.

The apparent contribution of individual barriers should also be interpreted in the context of the sequential deletion design. In our deletion series, Gabija was removed first, followed by Retron-I-B, Pf4 and the CTX-like prophage, before deletion of the type I restriction-modification system. Because type I restriction-modification produced such a large increase in susceptibility, differences among the preceding intermediate mutants were comparatively small in the full gradient and therefore more difficult to resolve. It is possible that Retron-I-B and resident prophages contribute subtle or phage-specific effects that would have been more apparent in a genetic background where type I restriction-modification had already been removed. Our results support a multilayered model in which some barriers have broad effects, while others make more context-dependent contributions to host permissiveness.

The different responses of the lytic and temperate phage panels further suggest that isolation history may shape apparent host range and susceptibility. Temperate phages in our panel displayed a strong permissiveness gradient across the deletion series, whereas many lytic phages showed more modest changes. Many of the lytic phages used here were originally isolated on PAO1 or other genetically unmodified hosts (Supplementary Table 4) and our lytic phage panel may therefore be enriched for phages that already infect such hosts efficiently. In contrast, our temperate phage collection was assembled using permissive host backgrounds, allowing recovery of phages that may be more strongly restricted by barriers present in MPAO1. Future experiments directly comparing newly isolated lytic phages recovered from matched environmental samples on MPAO1 versus viPAO1 would help disentangle potential effects of phage lifestyle from the effects of isolation host history.

Resident prophages are another important type of host-associated barrier. Prophages can protect bacterial hosts against superinfection through diverse mechanisms, including repressor-mediated immunity, surface modification, superinfection exclusion and intracellular defence functions [20]. Similar effects have also been observed in other *Pseudomonas* species, where resident cryptic prophages can provide strong protection against phage infection [24]. Removal of resident prophages from MPAO1 may therefore increase permissiveness both biologically, by eliminating prophage-mediated resistance, and practically, by reducing induction of resident prophages during phage propagation. It remains possible that some phages recovered using viPAO1 would be restricted in MPAO1 by immunity or superinfection exclusion encoded by resident prophages, particularly if they are closely related to resident MPAO1 prophages.

Surface receptor composition provided an additional route for altering phage recovery. Unlike removal of intracellular barriers, receptor-associated mutations did not simply make the host more permissive in a general sense. Instead, receptor mutants changed the composition of recovered phage populations. This likely reflects the role of receptors in shaping enrichment dynamics: abundant or rapidly replicating phages using a particular receptor can dominate mixed lysates and mask phages that use alternative infection routes. This interpretation is consistent with the broader understanding that host range is not binary, but reflects multiple stages of the infection cycle, including receptor recognition, adsorption, genome entry, intracellular replication and defence evasion, all of which can be influenced by host physiological state and assay conditions [1,25]. In our experiments, receptor-associated mutations had the clearest effect in MPAO1-derived backgrounds, suggesting that receptor-mediated shifts in recovery can be especially important when intracellular barriers still constrain which phages can propagate successfully.

By combining permissive and receptor mutant host backgrounds, we recovered a broad curated collection of temperate *P. aeruginosa* phages. This collection is a genomically diverse resource: the 69 individually purified phages represent 24 predicted genera, including 15 putative novel genera, and seven family-level groups, including one putative novel family-level group. Sequencing phages as lysogens further allowed confident identification of prophage boundaries and chromosomal insertion sites, linking each genome to a defined phage isolate. Importantly, this approach helps convert predicted prophage regions from bacterial genome data into experimentally tractable phage isolates. This is valuable because cultured, individually available phage collections remain essential for validating predictions from metagenomic and computational approaches, testing phage biology experimentally and exploring molecular mechanisms that cannot be resolved from sequence data alone.

Our findings also place host engineering alongside environmental sampling as a key axis for expanding cultured phage diversity. Many phage discovery studies attempt to increase diversity by sampling more locations, more environments or more bacterial isolates. These strategies are clearly valuable, and previous *Pseudomonas* phage isolation studies have shown that diverse phages can be recovered from environmental and clinical sources, particularly when multiple hosts are used [26–28]. However, our results show that sampling source is not the only determinant of recovered diversity. From the same induced lysates or environmental samples, different host backgrounds recovered different phages. Together with recent work in *E. coli*, where restoration of O-antigen and removal of antiviral barriers enabled recovery of previously inaccessible phage groups, our results support an emerging principle: engineering host permissiveness can reveal phage diversity that conventional hosts under-sample [29].

At the same time, permissive hosts reduce only one class of isolation bias. Even viPAO1 is unlikely to provide an unbiased view of natural phage diversity. Recovery still depends on enrichment conditions, culture medium, incubation temperature, host physiology, receptor compatibility, plaque formation, sequencing strategy and the ability of a phage to propagate under laboratory conditions. Environmental pooled sequencing also remains less resolved than individually purified phage isolation, particularly when closely related viruses are present in the same sample. More generally, it is unlikely that any single host background will capture the full diversity of phages associated with a bacterial species as genetically and phenotypically variable as *P. aeruginosa*. Future approaches may therefore benefit from combining engineered permissive hosts, receptor-modified derivatives, diverse clinical or environmental host backgrounds, and less selective isolation technologies.

Together, our findings demonstrate that intracellular defence systems, resident prophages and surface receptor composition collectively shape experimentally recoverable phage diversity in *P. aeruginosa*. By reducing these host-associated barriers, permissive host backgrounds enabled recovery of broad and previously under-sampled temperate phage diversity. Beyond phage isolation, viPAO1 may also provide a useful background for characterising individual phage-defence systems and defined prophage insertions while reducing confounding effects from endogenous defence systems and resident prophages. More broadly, this work supports the view that the host used for phage isolation shapes which phages we discover, and that host engineering should be considered alongside environmental sampling and methodological innovation as a strategy for uncovering hidden phage diversity.

## Materials and methods

### Bacterial strains and growth conditions

All bacterial strains used in this study are listed in Supplementary Table 1. *Escherichia coli* strains used for cloning and *Pseudomonas aeruginosa* strains were routinely cultured in Lysogeny Broth (LB) medium or on LB agar at 37°C. Liquid cultures were incubated with shaking at 180 rpm. Where appropriate, gentamicin was used at a final concentration of 20 µg ml^-1^ for *E. coli* and 50 µg ml^-1^ for *P. aeruginosa*.

The parental *P. aeruginosa* strain used for construction of viPAO1 was MPAO1. The complete MPAO1 genome has been previously described and is available under accession GCF_016107485.1 [30]. This genome differs from the PAO1 reference strain by several strain-specific regions, including an additional prophage region, making the MPAO1 reference particularly relevant for deletion design and downstream genome analysis.

*Pseudomonas aeruginosa* isolates used for prophage induction were obtained from six collections: a keratitis collection (n = 441; [reference/provider to be added]), the Thai clinical isolate collection (n = 57; [31]), the UMCU clinical isolate collection (n = 297; [reference/provider to be added]), a urinary tract infection collection (n = 62; [32,33]), cystic fibrosis clinical isolate collection (total n=511, 192 tested; [34]) and an international strain panel (n = 41; [35,36]). Together, these comprised 1090 isolates representing diverse clinical, environmental and reference backgrounds. Four isolates from this collection were selected for the controlled comparative induced-lysate isolation experiment described below. Collection-specific information, including isolate source and relevant references, is provided in Supplementary Table 2.

All strains were stored as glycerol stocks at −70°C. For routine experiments, strains were streaked from frozen stocks onto LB agar plates, and single colonies were used to inoculate overnight cultures.

### Construction of MPAO1 deletion mutants and viPAO1

Candidate anti-phage defence systems in MPAO1 were identified using DefenseFinder [11] and PADLOC [12] in June 2023, and resident prophage regions were predicted using PHASTEST [13] in June 2023. These analyses were used to define candidate defence and prophage regions for deletion.

The initial sequential deletion series was constructed in the laboratory MPAO1 stock available at the time. Deletions were introduced in the following order: Gabija, Retron-I-B, Pf4, the CTX-like prophage, the type I restriction-modification system and Helicase-DUF2290. The resulting strains were designated im1 to im5 and dPAO1 Δ*pilZ*. Specifically, im1 lacked Gabija; im2 additionally lacked Retron-I-B; im3 additionally lacked Pf4 prophage; im4 additionally lacked the CTX-like prophage; im5 additionally lacked the type I restriction-modification system; and dPAO1 Δ*pilZ* additionally lacked Helicase-DUF2290.

Whole-genome sequencing performed following construction of the initial deletion series showed that the parental MPAO1 stock carried an unintended mutation in *pilZ* and also revealed an additional resident prophage, Pf6, that had not been included in the original deletion design. The *pilZ* locus was subsequently restored to the MPAO1 reference allele by two-step allelic exchange in the parental MPAO1 strain and the corresponding im1–im5 and dPAO1 Δ*pilZ* derivatives. A repair construct carrying the reference *pilZ* allele together with flanking regions homologous to the MPAO1 chromosome was introduced using the same allelic-exchange procedure used for the deletion constructs, as described below. Candidate repaired clones were screened by PCR and confirmed by Sanger sequencing. The repaired dPAO1 strain was then used as the background for deletion of Pf6, generating the final virus-isolation strain viPAO1. Both the original dPAO1 Δ*pilZ* strain and the *pilZ*-repaired derivatives were retained and used in subsequent experiments as indicated in Supplementary Table 1 and the corresponding figure legends.

Unmarked deletions and restoration of the *pilZ* allele were generated by two-step allelic exchange using a *sacB*-based suicide-vector approach adapted from established methods for *P. aeruginosa* [37,38]. Upstream and downstream homology arms were amplified from the MPAO1 genome and ranged from approximately 500 to 2,000 bp depending on the size and structure of the target region. PCR products were assembled into the gentamicin-resistant suicide vector pDONORPex18GmR using NEBuilder HiFi DNA Assembly Master Mix. The resulting plasmids were introduced into *P. aeruginosa* by electroporation using 1-mm gap cuvettes and a Bio-Rad electroporator set to 1.8 kV. Single-crossover integrants were selected on LB agar containing gentamicin at 50 µg ml^-1^. Double-crossover mutants were selected on TYS10 agar containing 10% sucrose. Candidate sucrose-resistant colonies were screened for gentamicin sensitivity and tested by PCR across the expected deletion or allelic-replacement junction. PCR products from candidate mutants were confirmed by Sanger sequencing, and selected final engineered strains were subsequently verified by whole-genome sequencing.

Defence systems were deleted as single regions, whereas prophages were removed in one or more sections depending on their genomic structure: Pf4 was deleted in two steps, the CTX-like prophage in three steps and Pf6 in one step. Full strain genotypes and deletion coordinates are provided in Supplementary Tables 1 and 3c, respectively; primer sequences and plasmids used for strain construction are listed in Supplementary Tables 3a and 3b.

### Bacterial growth assays

Growth of MPAO1 and engineered deletion mutants was measured in LB medium using a 96-well plate-reader assay. Overnight cultures were diluted 1:100 into fresh LB and grown at 37°C to mid-exponential phase. Cultures were then diluted in fresh LB to a starting OD600 of 0.03, and 200 µl was transferred to each well of a 96-well plate.

Plates were incubated at 37°C for 18 h in a CLARIOstar Plus plate reader equipped with a stacker (BMG LABTECH). OD600 was measured every 30 min, with shaking before each measurement. Growth assays were performed with ten biological replicates per strain.

Maximum growth rate was calculated from each growth curve as the maximum increase in OD600 per hour between consecutive measurements using numpy.diff in Python.

### Phage collections and routine propagation

Phages used in this study are listed in Supplementary Tables 4 and 5. The lytic phage panel comprised 67 *Pseudomonas aeruginosa* phages obtained from previously described and collaborator-provided collections, including phages from the Citizen Phage Library [39], the collection used by Wright et al. [40], including phages originally described by Mattila et al. [41], and phages provided by Alexander Harms and collaborators [42]. Several CPL phages used in this study were additionally characterised by Tong et al. [43]. Other established *P. aeruginosa* phages in the panel have been described previously, including LUZ19 [44,45], LMA2 and 14-1 [46], UT1 [47], and PNM [48].

The temperate phage panel used for liquid infection assays comprised 43 *P. aeruginosa* phages isolated as part of the curated temperate phage collection described in this study, together with DMS3, which was already available in the laboratory. DMS3 was originally described by Budzik et al. [49].

For routine propagation, phages were recovered from frozen stocks and plaque-purified on permissive bacterial lawns where required. Individual plaques were picked and amplified overnight using viPAO1 grown in LB medium at 37°C with shaking. Following amplification, lysates were filtered through 0.22-µm filters to remove bacterial cells and debris.

Phage titres were determined by serial dilution using double-layer agar assays with LB bottom agar and 0.5% LB top agar. Phage lysates were stored in LB at 4°C for short-term use or as 25% glycerol stocks at −70°C for long-term storage.

### Liquid phage infection assays

Bacterial cultures were prepared as described for the growth assays. Briefly, overnight cultures were diluted 1:100 into fresh LB and grown at 37°C to mid-exponential phase, approximately OD600 0.6. Cultures were then diluted in fresh LB to OD600 0.03 and mixed with phage at a multiplicity of infection of 0.1. Each well contained a total volume of 200 µl. No-phage bacterial controls and LB blanks were included on each plate.

Plates were incubated at 37°C for 18 h in a CLARIOstar Plus plate reader equipped with a stacker (BMG LABTECH). OD600 was measured every 30 min, with shaking before each measurement. Infection assays were performed with at least three biological replicates per strain–phage combination.

Phage susceptibility was quantified from bacterial growth during liquid infection assays. Area under the OD600 growth curve (AUC) was calculated between 3 and 10 h post-infection by trapezoidal integration using numpy.trapz in Python. AUC values were normalised to the mean AUC of the viPAO1 no-phage control.

### Selection of prophage-rich clinical isolates

Clinical *P. aeruginosa* isolates from the keratitis collection were selected for the controlled comparative prophage induction experiment based on predicted prophage content. Genome assemblies generated as part of a parallel study were analysed using geNomAD v1.12.0 [50] to identify predicted prophage regions. Four isolates with the highest predicted prophage burden were selected for mitomycin C induction and comparative phage recovery across MPAO1, viPAO1 and derivative receptor-mutant host backgrounds. The four selected isolates and associated strain metadata are listed in Supplementary Table 1; sequencing and deposition information is provided in Supplementary Table 8.

### Generation of evolved receptor-mutant host derivatives

Evolved receptor-mutant derivatives of MPAO1 and viPAO1 were generated by selecting spontaneous phage-resistant mutants with altered surface-associated susceptibility. To select T4P mutants, MPAO1 and viPAO1 were challenged with phage CPL00211. To select LPS mutants, MPAO1 and viPAO1 were challenged with phage PA14P2.

Selection was performed using spot assays. Briefly, overnight cultures of MPAO1 or viPAO1 were mixed with LB soft agar and poured onto LB agar plates to generate bacterial lawns. Phage lysates were spotted onto the lawns and plates were incubated overnight at 37°C. Bacterial colonies arising within zones of lysis were picked, purified by streaking and screened for altered susceptibility to T4P- or LPS-dependent phages.

Candidate receptor-mutant derivatives were analysed by whole-genome sequencing to identify mutations associated with altered phage susceptibility. The evolved receptor-mutant hosts used in the comparative isolation experiments are listed in Supplementary Table 1, together with their corresponding genotypes. Sequencing and deposition information is provided in Supplementary Table 8.

### Whole-genome sequencing and genome analysis

The published MPAO1 genome sequence (GCF_016107485.1) was used as the reference for genome analysis.

Following construction of the deletion series, the parental MPAO1 strain used in this study, intermediate deletion mutants, dPAO1 and viPAO1 were sequenced to confirm the intended deletions (Supplementary Table 1). Short-read sequencing was performed by MicrobesNG (Birmingham, UK), including DNA extraction, library preparation and Illumina sequencing. In addition, viPAO1 was subjected to long-read sequencing to generate a complete genome assembly. Sequencing reads from engineered strains were analysed by comparison with the published MPAO1 reference genome. Sequencing datasets and associated deposition information are provided in Supplementary Table 8.

Four phage-evolved surface-receptor mutants used in the controlled comparative isolation experiments were also sequenced by MicrobesNG using Illumina short-read sequencing. Trimmed reads were analysed with breseq v0.40.1 by mapping to the corresponding parental genomes to identify genetic variants associated with the evolved phenotypes; de novo genome assemblies were not generated for these strains. Deposition information for these sequencing datasets is provided in Supplementary Table 8.

Four *P. aeruginosa* clinical isolates selected for the controlled comparative induced-lysate isolation experiment were sequenced using Oxford Nanopore long-read sequencing by Plasmidsaurus. Closed genome assemblies were obtained for all four isolates at approximately 80-90x sequencing coverage and were used to define their prophage content for the comparative isolation experiment. Sequencing and assembly deposition information is provided in Supplementary Table 8.

Genome assemblies for the broader *P. aeruginosa* isolate collection were generated as part of a parallel study and were used here to predict prophage content for strain selection. Assemblies were generated from MicrobesNG short-read sequencing data using a pipeline comprising FastQC v0.11.8, Trimmomatic v0.38, Shovill v0.9.0, QUAST v5.1.0rc1 and FastANI v1.33. Prophage content in these assemblies was predicted using geNomAD v1.12.0 [50].

### Comparative phage isolation from induced lysates and environmental samples

Phage recovery was compared across host backgrounds using two source types: induced prophage lysates from clinical *Pseudomonas aeruginosa* isolates and environmental samples. For the prophage induction experiment, four prophage-rich clinical isolates were selected based on predicted prophage content, as described above. Prophages were induced using mitomycin C following an adapted protocol previously described for *P. aeruginosa* [20]. Briefly, isolates were grown in LB at 37°C to an OD600 of approximately 0.5, and mitomycin C was added to a final concentration of 3 µg ml^-1^. Cultures were incubated until visible lysis was observed, approximately 4-5 h after induction. Induced cultures were centrifuged to remove bacterial debris, and supernatants were filtered through 0.22-µm filters to generate sterile induced lysates.

Environmental phages were isolated from four stored environmental samples: influent wastewater from Falmouth sewage treatment plant, influent wastewater from Carnon Downs sewage treatment plant, slurry from the Fraddon Biogas facility, and settled sludge from Marsh Mills wastewater treatment facilities. Sample collection procedures are described in Forsyth et al. [51]. Environmental samples were used directly for enrichment without prior centrifugation or filtration.

Both induced lysates and environmental samples were enriched on the same six host backgrounds: MPAO1, viPAO1, the MPAO1 T4P-associated mutant, the MPAO1 LPS-associated mutant, the viPAO1 T4P-associated mutant and the viPAO1 LPS-associated mutant. The generation and genotypes of these evolved receptor-mutant derivatives are described above and listed in Supplementary Table 1. For each enrichment, 5 ml LB was inoculated with 100 µl of overnight host culture and 100 µl of induced lysate or environmental sample, followed by overnight incubation at 37°C with shaking.

Following enrichment, lysates were filtered through 0.22-µm filters and plated on the corresponding enrichment host using double-layer plaque assays. Ten-fold serial dilutions were prepared in LB. For each dilution, 25 µl of diluted lysate was mixed with 150 µl of overnight host culture and 3 ml of 0.5% LB top agar and poured onto LB agar plates. Plates were incubated overnight at 37°C, and six well-separated plaques were picked from each sample-host combination.

Individual plaques were picked into 500 µl LB. To increase phage yield prior to pooled sequencing, 50 µl of each plaque suspension was used for a secondary enrichment on viPAO1. Following enrichment and filtration, lysates were pooled according to the original isolation host background. Equal volumes (150 µl) of individual lysates were combined for each host background, generating six pooled phage samples from the induced-lysate experiment and six pooled phage samples from the environmental experiment. Each pool contained up to 24 plaque-derived lysates, corresponding to six plaques from each of four input samples.

Phage DNA was extracted from pooled lysates using the Norgen Biotek phage DNA isolation kit according to the manufacturer’s instructions. Pooled phage DNA was subjected to Oxford Nanopore long-read sequencing by Plasmidsaurus. Sequencing and deposition information is provided in Supplementary Table 8. Assembly and recovery of phage genomes and contigs from the pooled sequencing data are described below.

### Isolation of the curated temperate phage collection

To isolate temperate phages at scale, 1090 *Pseudomonas aeruginosa* isolates from the five isolate collections described above were screened for inducible prophages (Supplementary Table 2).

Isolates were induced with mitomycin C using the conditions described above. Inductions were performed in 96-well deep-well plates, and induced lysates were filtered using 96-well filter plates. To identify lysates containing phages capable of infecting the principal isolation hosts, undiluted induced lysates were spotted onto lawns of viPAO1, dPAO1 and dPAO1 Δ*pilZ*. dPAO1 Δ*pilZ* was included to reduce repeated recovery of T4P-dependent transposable phages and facilitate recovery of additional phage types.

Lysates producing visible lysis on at least one isolation host were selected for plaque purification. Serial dilutions of positive lysates were plated on the corresponding permissive host using double-layer plaque assays, and individual well-separated plaques were picked. Each phage isolate was plaque-purified three times using its original isolation host (viPAO1, dPAO1 or dPAO1 Δ*pilZ*). Purified phages were amplified on permissive hosts, filtered through 0.22-µm filters and stored as described above.

To generate lysogens for genome sequencing and downstream characterisation, purified temperate phages were plated on dPAO1, viPAO1 or dPAO1 Δ*pilZ*, and resistant colonies arising from turbid plaques or lysis zones were streak-purified. Candidate lysogens were screened for resistance to superinfection by the corresponding phage and induced with mitomycin C to confirm production of infectious phage particles. Confirmed lysogens were stored as glycerol stocks at −70°C and processed for whole-genome sequencing as described below.

Using this workflow, 78 plaque-purified phage isolates were recovered, of which 75 formed stable lysogens under the conditions tested. Following recovery of phage genomes from lysogen sequencing, isolates were compared at the genome level to identify repeated recovery of genetically identical phages. Independently recovered isolates with identical genome sequences were retained as separate records in the collection metadata, particularly when recovered from different source isolates, and were distinguished using numerical suffixes where appropriate. For genome-level analyses, however, identical genomes were collapsed to a single representative.

Among phages recovered using the PAO1-derived isolation hosts, this dereplication yielded 71 distinct genome groups. Two of these corresponded to independent re-isolations of the previously described phages JBD24 and JBD69 and were therefore excluded from the count of newly recovered phage diversity. The principal curated collection used for collection-level diversity analyses consequently comprised 69 unique newly isolated temperate phages. Phage isolate metadata, including source isolate, isolation host, lysogen host and genome-level dereplication group, are provided in Supplementary Table 5.

Phage names were assigned using a consistent naming scheme in which “PA” denotes *Pseudomonas aeruginosa* and is incorporated into a recognisable word or phrase. These names are used as isolate identifiers only and do not imply taxonomic relatedness, receptor usage or phenotypic similarity. Where the same phage genome was recovered independently on multiple occasions, the individual isolates were retained and distinguished using numerical suffixes.

Three phages in the principal curated collection, PAndora, PAranormal and PAulmccartney, formed plaques on the isolation hosts but did not form stable lysogens in dPAO1 or viPAO1 under the conditions tested. These phages were therefore sequenced directly from phage lysates rather than recovered from lysogen sequencing and were retained in the 69-phage curated collection.

The same prophage-induction, plaque-purification and lysogen-validation workflow was also applied using *P. aeruginosa* PA14 [52] as an isolation host. This yielded 12 independently recovered phage isolates representing 11 unique phage genomes, (with PAneer1 and PAneer2 being independent isolates with identical genome sequences). Nine of the PA14-isolated phages were also capable of infecting PAO1. These phages are included in Supplementary Table 5 and deposited alongside the broader collection but were excluded from the principal 69-phage collection and from the corresponding collection-level diversity analyses.

### Phage genome sequencing, annotation and prophage boundary identification

For the curated temperate phage collection, phage genomes were recovered from whole-genome sequencing of lysogens. Between one and four independently generated lysogens were sequenced per phage, with two lysogens sequenced for most phages. Long-read whole-genome sequencing and assembly of bacterial lysogens was performed by MicrobesNG.

Lysogen genome assemblies were analysed to identify prophage regions corresponding to the infecting phage. Predicted viral regions were identified using geNomAD v1.12.0 [50] and compared against the corresponding uninfected host genome, dPAO1, dPAO1 dPilZ or PA14, to define prophage boundaries. Prophage boundaries and chromosomal insertion sites were manually curated by comparison with the corresponding uninfected host genome. For each lysogen, sequences spanning the left and right prophage-chromosome junctions were compared with the corresponding chromosomal sequence prior to phage insertion. Shared sequence motifs present at both prophage junctions and corresponding to the uninterrupted chromosomal locus were used to define the attachment sites (attB) sequences. Where multiple independently generated lysogens were available for the same phage, junction sequences and insertion sites were compared to confirm consistency (Supplementary table 5).

Predicted phage genomes were annotated using Pharokka v1.7.3 [53]. Final genome sequences and annotations are provided under accession numbers listed in Supplementary Tables 5 and 8.

For three phages recovered during the screening that formed plaques but did not form stable lysogens in dPAO1 or viPAO1 under the conditions tested, phage DNA was extracted directly from lysates using the Norgen Biotek phage DNA extraction kit, according to the manufacturer’s instructions. DNA from these lysates was sequenced using long-read sequencing by Plasmidsaurus. Oxford Nanopore reads were quality assessed with NanoPlot v1.46.2 and filtered with Chopper v0.11.0, retaining reads with a minimum quality score of 10 and minimum length of 1,000 bp. Residual host-derived reads were removed by mapping filtered reads to the viPAO1 host genome with minimap2 v2.30 [54] using the ONT preset; unmapped reads were retained using samtools v1.23.1. Assemblies were generated with Flye v2.9.6 [55] and polished with Medaka v2.2.1. Resulting phage genomes were annotated as described above.

### Pooled phage sequencing and genome recovery

Pooled phage DNA samples from the comparative induced-lysate and environmental isolation experiments were sequenced using long-read sequencing by Plasmidsaurus. Raw reads were quality-checked with NanoPlot v1.46.2. Reads were filtered with chopper v0.12.0 using a minimum read quality of 10 and a minimum read length of 1,000 bp. Because all plaque-derived lysates were secondarily enriched on viPAO1 before pooling, filtered reads were mapped to the viPAO1 host genome with minimap2 v2.30 using the map-ont preset. Reads that did not align to viPAO1 were extracted with samtools v1.23.1 and used as the host-depleted read set for downstream analyses.

For environmental samples, host-depleted reads were assembled *de novo*. Assemblies were generated with Flye v2.9.6 in metagenome mode using --meta and the Nanopore high-quality read model (--nano-hq) [56]. Contigs were polished with Medaka v2.2.1. Candidate contigs of at least 10 kb were retained for viral annotation and quality assessment.

Polished environmental assemblies were analysed with geNomAD v1.12.0 and CheckV v1.0.3 using the end-to-end workflow in both cases. Contigs were manually curated to retain medium-to-high confidence phage-derived contigs, defined as contigs with a geNomAD viral score greater than 0.9 and CheckV-estimated phage genome completeness of at least 50%. Retained contigs were compared all-versus-all with minimap2 v2.30, and near-duplicate contigs were clustered when pairwise identity was at least 90% and alignment coverage of the shorter contig was at least 50%. One representative per cluster was selected, favouring the longest contig. This generated a curated catalogue of 28 phage contigs recovered from the environmental pooled sequencing samples.

For induced-lysate pools, host-depleted reads were mapped to the full bacterial reference genomes corresponding to the four clinical isolates from which prophages had been induced. Mapping was performed with minimap2 v2.30. Candidate prophage-enriched regions were identified from primary-alignment depth using reads with mapping quality of at least 20. Regions of at least 10 kb with read depth of at least 5 were retained as candidate induced temperate phage genomes. This generated a catalogue of 9 induced phage genomes recovered across the induced-lysate pooled sequencing samples.

To determine which phages or phage contigs were present in each pooled sample, viPAO1-depleted reads from induced-lysate and environmental samples were mapped back to the corresponding curated catalogue using minimap2 v2.30 with secondary alignments suppressed. A phage genome or phage contig was considered present in a sample if primary alignments with MAPQ ≥20 covered at least 50% of its length or 10 kb, whichever was greater, and if at least five reads aligned to the genome or contig.

### Phage taxonomy and gene-sharing network analysis

Phage taxonomy and gene-sharing relationships were analysed using vConTACT3 [16] with a reference dataset derived from the vConTACT3 database. Reference genomes were filtered to include *Pseudomonas aeruginosa* phages only. Two separate vConTACT3 analyses were performed: one including the 69 individually purified temperate phages from the curated collection, and one including phage genomes and phage contigs recovered from the comparative induced-lysate and environmental isolation experiments.

Predicted family- and genus-level taxonomic assignments were obtained from vConTACT3 output. Phages assigned to vConTACT3 clusters containing reference phages with existing ICTV taxonomy were assigned to the corresponding taxonomic group where supported by the network structure. Phages belonging to vConTACT3 clusters that lacked reference phages with ICTV genus-level taxonomy were considered putative novel genera. Phages for which vConTACT3 did not identify a cluster containing reference phages with ICTV family-level taxonomy were considered to belong to a putative novel family.

Gene-sharing networks generated by vConTACT3 were visualised in Cytoscape v3.10.4 [57]. Network layouts were retained as generated, without manual repositioning of nodes. For the comparative isolation experiments, vConTACT3 assignments were used to compare the diversity of phages and phage contigs recovered on different host backgrounds.

## Statistical analysis

Statistical analyses were performed in Python 3.10.14. For bacterial growth assays, maximum observed growth rate was calculated for each biological replicate as the maximum increase in OD600 per hour between consecutive measurements. Differences between strains were tested using two-sided paired t-tests across matched experimental runs, and P values from multiple pairwise comparisons were adjusted using the Benjamini-Hochberg false-discovery-rate correction.

For liquid phage infection assays, area under the OD600 growth curve (AUC) between 3 and 10 h post-infection was used as the response variable. For each phage, differences in normalised AUC between host backgrounds and viPAO1 were tested using a least-squares linear model with normalised AUC as the now response variable and strain as a categorical predictor. viPAO1 was specified as the reference strain. P values were adjusted for multiple testing using the Benjamini–Hochberg method. Statistical significance was defined as an adjusted P value < 0.05.

## Data availability

Genome assemblies and sequencing reads generated in this study will be deposited in NCBI under several BioProject accessions (Supplementary table 8). Individual accession numbers for bacterial genomes, phage genomes and pooled sequencing datasets are provided in Supplementary Tables 1, 5, 8. The complete viPAO1 genome assembly will be deposited under accession [XXXX]. Genome sequences for the curated temperate phage collection, including phage name, source isolate, isolation host, lysogen host, insertion site and accession number, are listed in Supplementary Table 5.

Sequencing data generated from pooled induced-lysate and environmental phage samples will be deposited under accession numbers [XXXX-XXXX]. Curated phage genomes and phage contigs recovered from these pooled sequencing datasets are provided in Supplementary Tables 6 and 7. Genome sequences for the three non-lysogen-forming phages sequenced directly from lysates are also listed in Supplementary Table 5.

Previously published phage and bacterial genome sequences used in this study were obtained from public databases, and accession numbers are provided in Supplementary Tables 1 (strains), 4 (phages).

The code used for the analyses described in this study is available at https://github.com/AlekseiAgapov/viPAO1.

---

**Supplementary Figure 1. Expanded phage-susceptibility and growth profiling of the MPAO1 deletion series.**

(A,B) Susceptibility of intermediate deletion strains derived from *P. aeruginosa* MPAO1 to full panels of lytic phages (A) and temperate phages (B). The deletion series comprises intermediate mutants (im1–im5), dPAO1 and viPAO1, as shown in Fig. 1A. Heatmap values were derived from growth-based infection assays performed in LB at 37°C using an MOI of 0.1. Bacterial growth in the presence or absence of phage was monitored by OD_600_ time-course measurements, and growth was summarised as the area under the growth curve (AUC) between 3 and 10 h. For each strain-phage combination, AUC values from independent experimental runs were pooled and normalised to the mean AUC of the viPAO1 no-phage control. Lower normalised AUC values indicate stronger phage-mediated growth inhibition and therefore greater susceptibility. Asterisks indicate strain-phage combinations with significantly higher normalised AUC values relative to viPAO1 following Benjamini-Hochberg correction for multiple testing (adjusted P < 0.05). Most strain–phage combinations were tested across five independent experiments, while no-phage controls were included in ten independent experiments.

(C) Maximum growth rates of the MPAO1 deletion series grown in LB at 37°C under standard laboratory conditions. Growth rates were calculated from OD600 growth curves obtained in plate-reader assays. Points represent independent biological replicates (n = 10); black points and error bars indicate mean ± SEM.

**Supplementary Figure 2. Host genotype shapes recovery of environmental phage contigs.**

(A) Schematic overview of the comparative environmental phage isolation workflow. Four environmental samples were enriched on six host backgrounds: MPAO1, viPAO1, and corresponding T4P- and LPS-associated receptor-mutant derivatives. Following plaque assays, six plaques were picked from each sample-host pair. Plaque-derived lysates were secondarily enriched on viPAO1, then pooled by original isolation host background to generate six host-specific sequencing pools. Pooled phage DNA was extracted and sequenced using long-read sequencing.

(B) Presence/absence matrix of environmental phage contigs detected in host-specific sequencing pools. Columns represent curated environmental phage contigs with a geNomAD viral score > 0.9 and CheckV-estimated completeness of at least 50%; rows represent the host backgrounds used for phage isolation. A phage contig was considered present in a sequencing pool if primary alignments with MAPQ ≥ 20 covered at least 50% of the contig length or 10 kb, whichever was greater, and at least five reads aligned to the contig.

(C) Genus-level summary of environmental phage recovery across host backgrounds based on vConTACT3 assignments.

**Supplementary Figure 3. Comparative isolation recovers phages across diverse gene-sharing groups.**

vConTACT3 gene-sharing network comparing the nine induced temperate phage genomes and 28 environmental phage contigs recovered in the comparative isolation experiments with publicly available *P. aeruginosa* reference phages. Nodes represent phage genomes or contigs and edges represent gene-sharing relationships inferred by vConTACT3. Environmental phage contigs are shown in green, phages recovered from induced lysates in orange, and reference phages in grey. Labels identify individual recovered phages and contigs using the same IDs as in Fig. 2 and Supplementary Fig. 2. Insets enlarge selected regions containing recovered sequences. Recovered induced phages and environmental contigs are distributed across multiple regions of the network, indicating that host-dependent recovery extends across genomically distinct phage groups.

**Supplementary Figure 4. Taxonomic and genome-size diversity of the curated temperate phage collection.**

(A) Taxonomic composition of the 69 unique newly isolated temperate phages included in collection-level analyses. Phages were classified using vConTACT3 into 24 predicted genera distributed across seven predicted families. The collection includes 15 putative novel genera and one putative novel family; established and putative novel family- and genus-level assignments are indicated separately.

(B) Distribution of phage genome lengths across predicted family-level groups represented in the curated collection. Individual points represent phage genomes, with boxplots summarising the genome-length distribution within each family.

**Supplementary Table 1. Bacterial strains used in this study.**

Bacterial strains used for strain construction, phage isolation, propagation, lysogen generation and comparative infection experiments. For each strain, the parental background, genotype or genetic modification, relevant phenotype or experimental use, construction or origin, source/reference and genome accession are provided. The MPAO1-derived deletion series includes strains carrying the original unintended *pilZ* mutation and corresponding derivatives in which the *pilZ* reference allele was restored. viPAO1 represents the final engineered virus-isolation host following removal of the identified intracellular defence systems and resident prophages.

**Supplementary Table 2. *Pseudomonas aeruginosa* isolate collections screened for inducible phages.**

Clinical and reference *P. aeruginosa* isolate collections screened for inducible prophages in this study. The table lists the number of isolates screened from each collection, infection or source type, relevant reference or provider, and availability of genomic data. A total of 1090 isolates were screened.

**Supplementary Table 3a. Primers used in this study.**

Oligonucleotide primers used for construction and verification of engineered *P. aeruginosa* strains. Primer sequences are shown in the 5′–3′ orientation, together with their principal experimental application.

**Supplementary Table 3b. Plasmids used in this study.**

Plasmids used for allelic exchange and construction of the MPAO1-derived deletion series and *pilZ*-repaired strains. The table provides plasmid name, description and source or reference.

**Supplementary Table 3c. Genomic regions deleted during construction of the MPAO1-derived permissive host series.**

Defence-system and prophage regions removed during construction of the engineered MPAO1 derivatives. Coordinates refer to the indicated MPAO1 genome assembly and specify the regions deleted for Gabija, Retron-I-B, Pf4, the CTX-like prophage, type I restriction–modification, Helicase-DUF2290 and Pf6.

**Supplementary Table 4. Lytic phage panel used for susceptibility assays.**

Lytic *P. aeruginosa* phages used to assess susceptibility across the MPAO1-derived deletion series. For each phage, the original isolation host where known, isolation source, provider, reference and genome accession are listed.

**Supplementary Table 5. Temperate phages recovered and characterised in this study.**

Temperate phages recovered following prophage induction and plaque isolation using PAO1-derived permissive hosts and PA14. The table provides source isolate and collection, isolation host, genome length, known or inferred receptor class, chromosomal insertion locus or loci, predicted bacterial attachment site (attB), inferred integration mode, and vConTACT3 family-, genus- and order-level assignments. Independently recovered isolates with identical genome sequences are retained as separate records and distinguished by numerical suffixes. For the principal collection-level analyses, duplicate genome recoveries were collapsed to a single representative and isolates identical to the previously described phages JBD24 and JBD69 were excluded, yielding 69 unique newly isolated phages recovered using PAO1-derived hosts. In addition, the table includes 12 phage isolates recovered using PA14 as the isolation host; these were retained in the collection metadata but excluded from the principal 69-phage analyses. Putative novel family- and genus-level assignments are indicated.

**Supplementary Table 6. Phage genomes recovered from the comparative induced-lysate isolation experiment.**

Nine unique induced temperate phage genomes recovered following enrichment and plaque isolation on MPAO1, viPAO1 and their T4P- and LPS-associated receptor-mutant derivatives. Host-specific presence/absence calls are shown for each phage genome. A genome was considered detected in a sequencing pool if primary alignments with MAPQ ≥ 20 covered at least 50% of the genome length or 10 kb, whichever was greater, and at least five reads aligned to the genome. Genome length, sequencing-support metrics and vConTACT3 genus- and family-level assignments are also provided.

**Supplementary Table 7. Environmental phage contigs recovered in the comparative isolation experiment.**

Twenty-eight unique environmental phage contigs recovered following enrichment and plaque isolation from four environmental samples on MPAO1, viPAO1 and their T4P- and LPS-associated receptor-mutant derivatives. Host-specific presence/absence calls are shown for each contig. Candidate phage contigs were retained following geNomAD and CheckV quality assessment and dereplication as described in Methods. A contig was considered detected in a host-specific sequencing pool if primary alignments with MAPQ ≥ 20 covered at least 50% of the contig length or 10 kb, whichever was greater, and at least five reads aligned to the contig. The table also provides contig length, geNomAD viral score, CheckV completeness and quality category, read-mapping support, and vConTACT3 taxonomic assignments.

**Supplementary Table 8. Sequencing datasets and accession information.**

Summary of sequencing datasets generated in this study, including engineered bacterial strains, receptor-mutant hosts, single-prophage lysogens, individually recovered phage genomes, pooled comparative-isolation libraries and the clinical isolates used as prophage sources. For each dataset group, the number of samples or genomes, deposited data type, sequencing platform/provider and relevant BioProject, BioSample, SRA and assembly/GenBank accessions are provided.

## Author contributions

**Anna Olina:** Conceptualization, Investigation, Methodology, Visualization, Writing – original draft, Writing – review & editing.

**Aleksei Agapov:** Conceptualization, Formal analysis, Data curation, Software, Visualization, Funding acquisition, Writing – review & editing.

**Xinyan Yu:** Investigation, Funding acquisition, Writing – review & editing.

**Rama P. Bhatia:** Resources, Writing – review & editing.

**Joanne L. Fothergill:** Conceptualization, Resources, Funding acquisition, Writing – review & editing.

**Michael A. Brockhurst:** Conceptualization, Resources, Funding acquisition, Writing – review & editing.

**Stineke van Houte:** Conceptualization, Supervision, Funding acquisition, Writing – review & editing.

**Edze R. Westra:** Conceptualization, Supervision, Project administration, Funding acquisition, Writing – review & editing.

## Acknowledgements

We thank Rosanna Wright, Rob Lavigne and Alexander Harms for providing phages used in this study. We thank Ben Temperton and the Citizen Phage Library, in particular Ellie Tong, Julie Fletcher and Christian Fitch, for providing phages and associated resources. We thank Sean Meaden for helpful discussions regarding phage sequencing and experimental design, and James Hall for valuable early discussions that helped shape the direction of the research. We thank Angus Buckling for providing the original MPAO1 strain and Marcel de Zoete (UMC Utrecht) for providing the Utrecht *Pseudomonas aeruginosa* strain collection.

