## Supplementary figures and images for "Engineering the Pseudomonas aeruginosa virus-isolation host viPAO1 reveals how host genetic barriers shape recoverable phage diversity"

### Supplementary figure 1

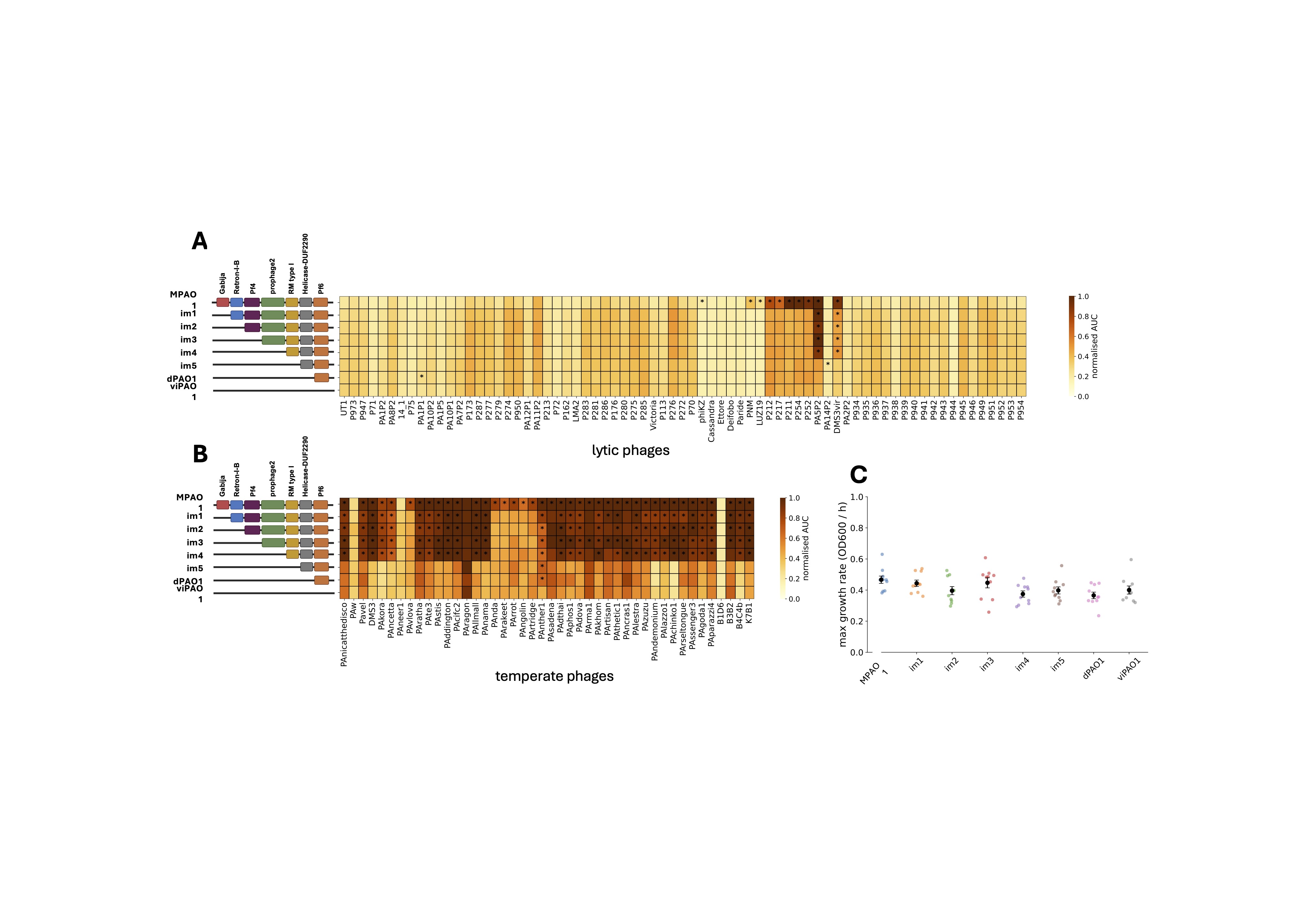

### Supplementary figure 2

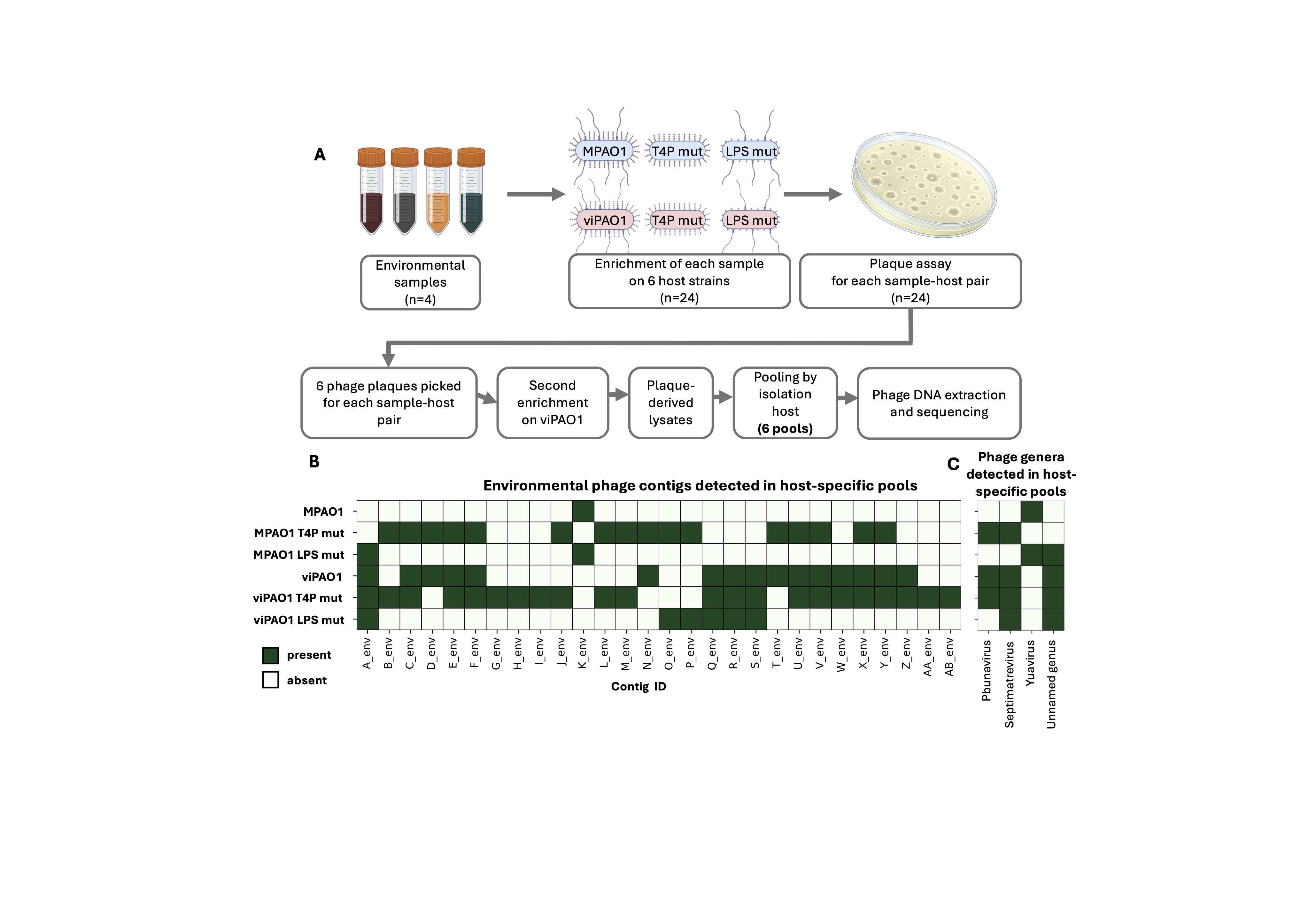

### Supplementary figure 3

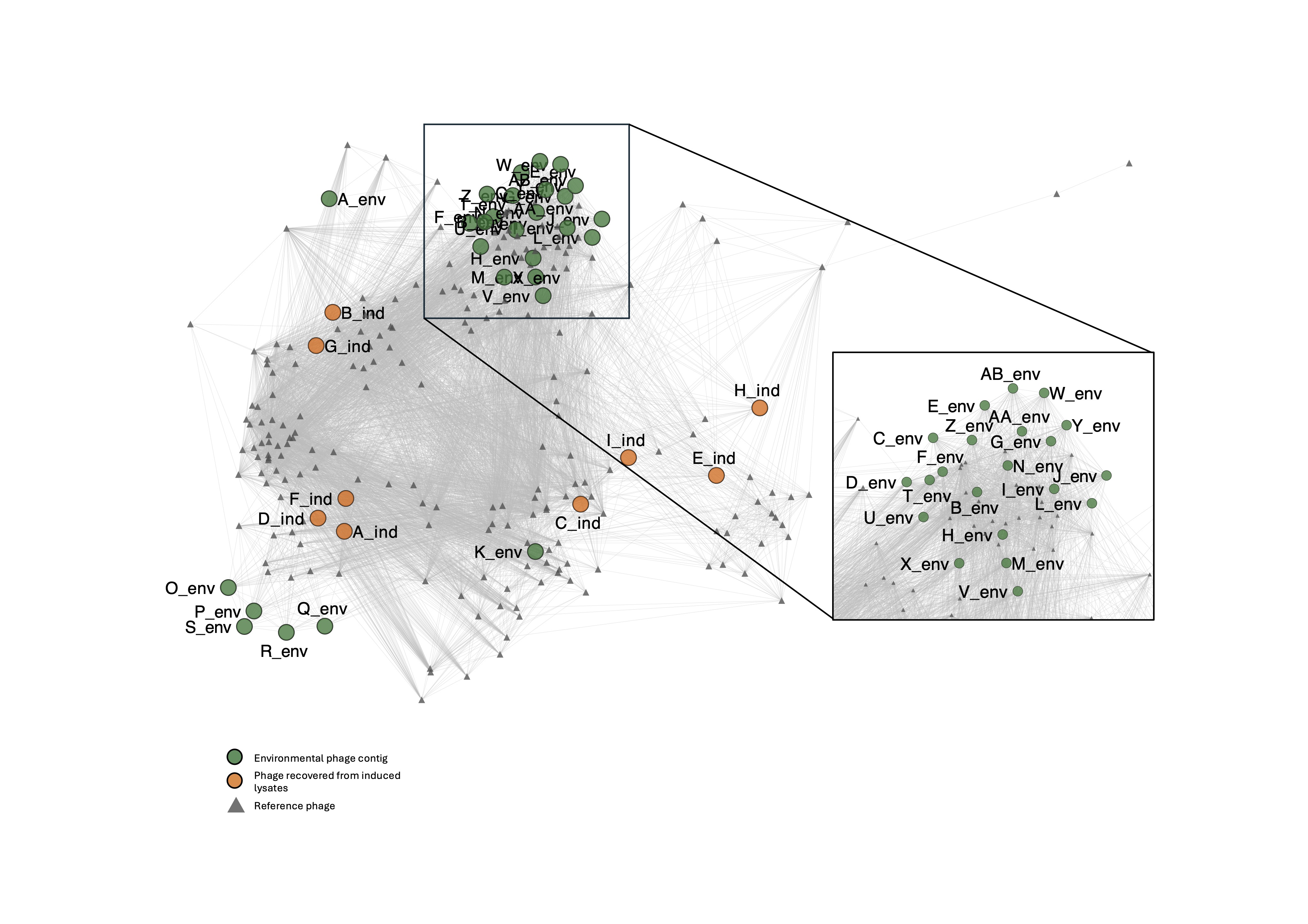

### Supplementary figure 4

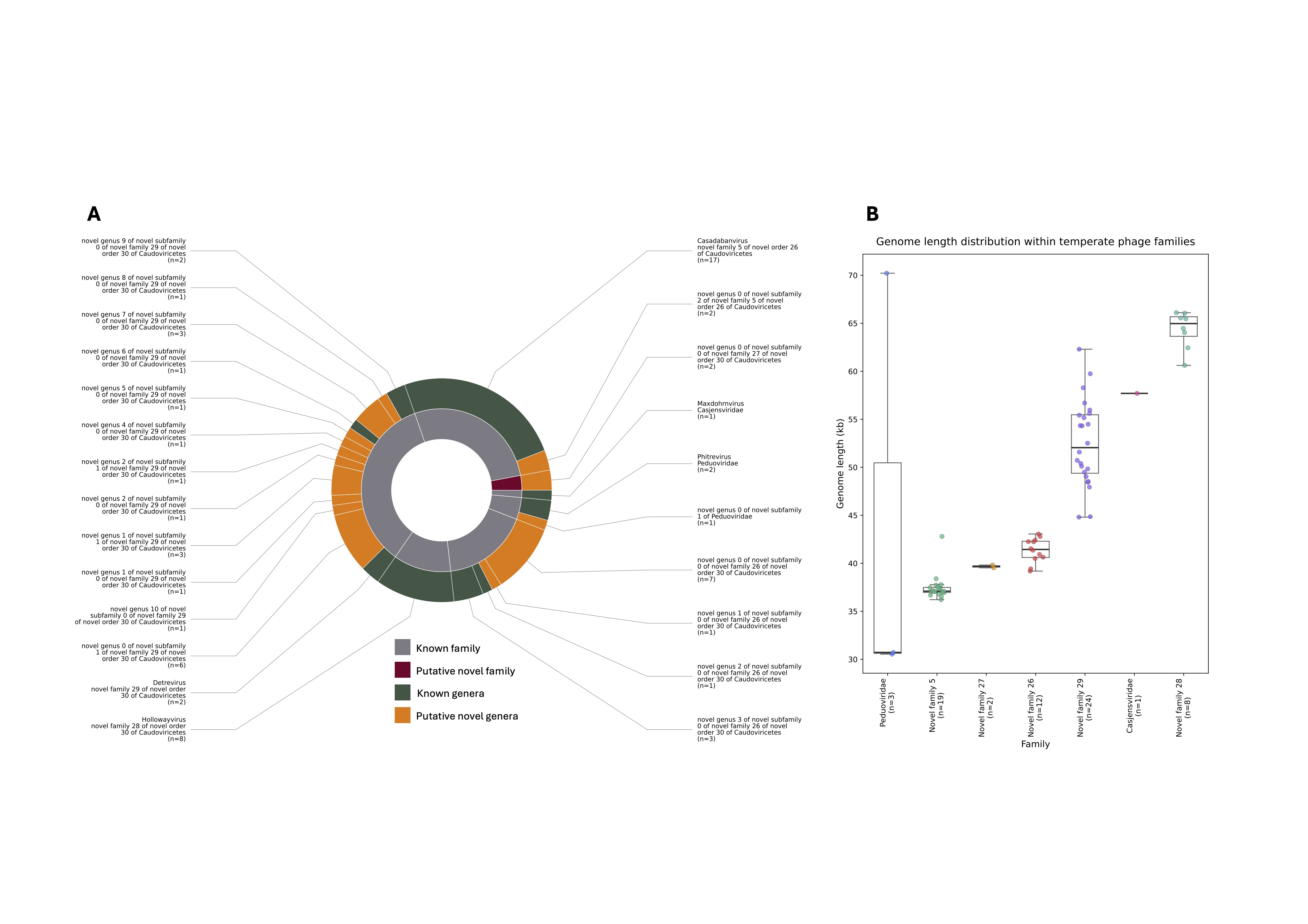
